# *Ex vivo* osteochondral test system with control over cartilage defect depth – a pilot study to investigate the effect of oxygen tension and chondrocyte-based treatments in chondral and full thickness defects in an organ model

**DOI:** 10.1101/2020.11.06.348177

**Authors:** Andrea Schwab, Alexa Buss, Oliver Pullig, Franziska Ehlicke

## Abstract

**Objective:** Cartilage defect treatment strategies are dependent on the lesion size and severity. Osteochondral explants models are a platform to test cartilage repair strategies *ex vivo*. Current models lack in mimicking the variety of clinically relevant defect scenarios. In this controlled laboratory study, an automated device (artificial tissue cutter, ARTcut^®^) was implemented to reproducible create cartilage defects with controlled depth. In a pilot study, the effect of cartilage defect depth and oxygen tension on cartilage repair was investigated.

**Design:** Osteochondral explants were isolated from porcine condyles. 4 mm chondral and full thickness defects were treated with either porcine chondrocytes (CHON) or co-culture of 20 % CHON and 80 % MSC (MIX) embedded in collagen hydrogel. Explants were cultured with tissue specific media (without TGF-β) under normoxia (20 % O_2_) and physiological hypoxia (2 % O_2_). After 28 days, immune-histological stainings (Collagen II and X, Aggrecan) were scored (modified Bern-score, 3 independent scorer) to quantitatively compare treatments outcome.

**Results:** ARTcut^®^ represents a software-controlled device for creation of uniform cartilage defects. Comparing the scoring results of the MIX and the CHON treatment, a positive relation between oxygen tension and defect depth was observed. Low oxygen tension stimulated cartilaginous matrix deposition in MIX group in chondral defects and CHON treatment in full thickness defects.

**Conclusion:** ARTcut^®^ has proved a powerful tool to create cartilage defects and thus opens a wide range of novel applications of the osteochondral model, including the relation between oxygen tension and defect depth on cartilage repair.

## 1 Introduction

Defects of the articulating surface are frequently occuring diseases in the field of orthopedics, traumatology and sports medicine [1]. Around 14 % of patients who suffered from a trauma induced knee injury during their middle ages, including fractures of the tibia, fibula, femur, or patella, develop osteoarthritis in the same knee joint at later age (>65 years) [2]. High risk of failure of more than 20 % in cartilage defect repair require to rethink and improve current cartilage treatment concepts with the aim for long-term repair of cartilage defects [3, 4].

Due to the avascular and aneural character of adult articular cartilage, large defects do not heal spontaneously [5]. Defects that are classified according to International Cartilage Repair Society (ICRS) grade III (defect depth >50 % cartilage depth) and IV (defect reaches subchondral bone) need surgical intervention to repair [6]. Gold standard for defects larger than 2-3 cm^2^ is the autologous chondrocyte implantation (ACI), resulting in similar outcome as microfracture [7, 8]. In a recent data analysis for cartilage defects, matrix-associated chondrocyte implantation (M-ACI) showed a significant lower reoperation rate than microfracture 2 years post-op [9]. M-ACI as well as ACI require two surgical interventions; the first to take a cartilage tissue sample for subsequent chondrocyte isolation and the second one for injection of isolated *in vitro* expanded autologous chondrocytes into defect site.

However, (M-)ACI has two main limitations: Due to the limited number of healthy and non-degenerated chondrocytes, cells have to be expanded *in vitro* to achieve sufficient number of cells needed for the implantation (ACI: 1 million cells per cm^2^, M-ACI: 20 million cells per cm^3^) [10–12]. Further, chondrocyte-based cartilage treatments, as well as microfracture techniques, result in formation of fibrocartilage, characterized by its inferior mechanical properties and thus lacking functional restoration compared to healthy hyaline cartilage [4, 13–17]. One approach to overcome the limitation of cell number for (M-)ACI treatment are MSC-chondrocyte co-cultures, *e.g*., 80 % MSC and 20 % chondrocytes, that reduces autologous chondrocyte cell number at same total cell density. Co-cultures of MSCs and chondrocytes have shown to increase cartilaginous matrix production and reduce hypertrophy, associated with MSCs during chondrogenic differentiation [18, 19].

To bring cartilage treatments to the next level, cartilage repair needs to be studied and understood in more detail, starting with basic research questions in a physiologically relevant environment.

An *ex vivo* cartilage defect test system based on osteochondral explants was established by Schwab *et al.* [20]. This test system represents a valuable *ex vivo* platform for biomaterial evaluation in critical size trauma-induced cartilage defects in terms of biocompatibility, biomaterial tissue integration and cartilage repair with higher throughput compared to *in vivo* models. Separated media compartments of the culture device allow for controlled, tissue- and cell-specific nutrient supply during *ex vivo* culture of osteochondral explants. This model also allows direct comparison of different treatment approaches under controlled conditions and has been shown to stimulate cartilage-like tissue formation of chondrocytes or MSCs in osteochondral lesions [21, 22].

However, this defect model was limited to full thickness defects created with a biopsy punch. To date, the creation of defects that do not fully penetrate the cartilage with the help of a biopsy punch or scalpel suffer from low reproducibility and are dependent on the operator. Therefore, an automated device with control over drilling depth to create defects with high reproducibility can overcome this limitation to study chondral wound healing. Of note, the drilling should not harm surrounding cartilage tissue and cell viability, neither by mechanical tissue disruption nor by friction induced heating. For the creation of defects in hard tissues like cartilage or even bone, there is no device available for automated defect creation that meets the above-mentioned requirements.

Following, in the present study the defect creation of the *ex vivo* defect model introduced by Schwab *et al*. was modified by implementation of a semi-automated drilling device, originally developed to mechanically induce standardized and reproducible wounds in full thickness skin equivalents [23]. Key features of this artificial tissue cutter (ARTcut^®^) are the sensor controlled optical barrier in combination with a moveable milling machine along x-, y- and z-axis. The light-barrier controlled drilling allows for creation of defects with defined depths in a reproducible set-up. Creation of tissue defects with ARTcut^®^ can be performed under sterile conditions for subsequent *in vitro* or *ex vivo* culture with the possibility to adjust defect geometry and control of defect creation process.

Another fact that has been neglected in the studies by Schwab *et al.* is the low oxygen tension present in the articular capsule [24]. It has been shown in literature that low oxygen tension is an important stimulus in chondrogenesis and increases the chondrogenic potential of articular cartilage progenitor cells and MSCs [24–26]. Taken together, the present study aimed to implement the ARTcut^®^ as tool for automated wounding to the *ex vivo* model allowing to study the influence of defect depth (full thickness defects and chondral defects) on tissue repair. In a pilot study, the relation between oxygen tension (normoxia 20 % vs. physiological hypoxia 2 %), defect depth and treatment strategy was studied. For the treatment strategies either chondrocytes only (CHON) or a mixture of 80 % MSC and 20 % CHON (MIX) were embedded in a collagen type I hydrogel for implantation into cartilage defects in the *ex vivo* osteochondral model without further supplementation of growth factors to elusively study the effect of the culture conditions on cartilage repair.

## 2 Methods

### 2.1 Osteochondral cylinders: Isolation, defect creation and *ex vivo* culture

Osteochondral cylinders (diameter: 8 mm, height: 5 mm) were isolated from medial femoral condyles of 6-8-month-old domestic pigs as previously described [20]. Creation of defects in osteochondral cylinders was carried out either manually or automatically, depending on the intended defect depth: Full thickness defects were manually created with a biopsy punch (diameter 4 mm; Kai Medical, BPP-40F). Chondral defects of 1 mm depth were created using custom-built ARTcut^®^ (Figure 1A), a software-controlled device for automatic wound placement in tissues developed by Fraunhofer ISC and IGB, Wuerzburg [23]. Osteochondral cylinders were fixed in support plate (Figure 1C) at specific x- and y-positions, and onset of drilling 1 mm chondral defects (z-axis) was determined with light beam (Figure 1B). For more details on the standardized defect creation process using ARTcut^®^, see chapter 1.2 in the supplementary information.

**Figure 1:**
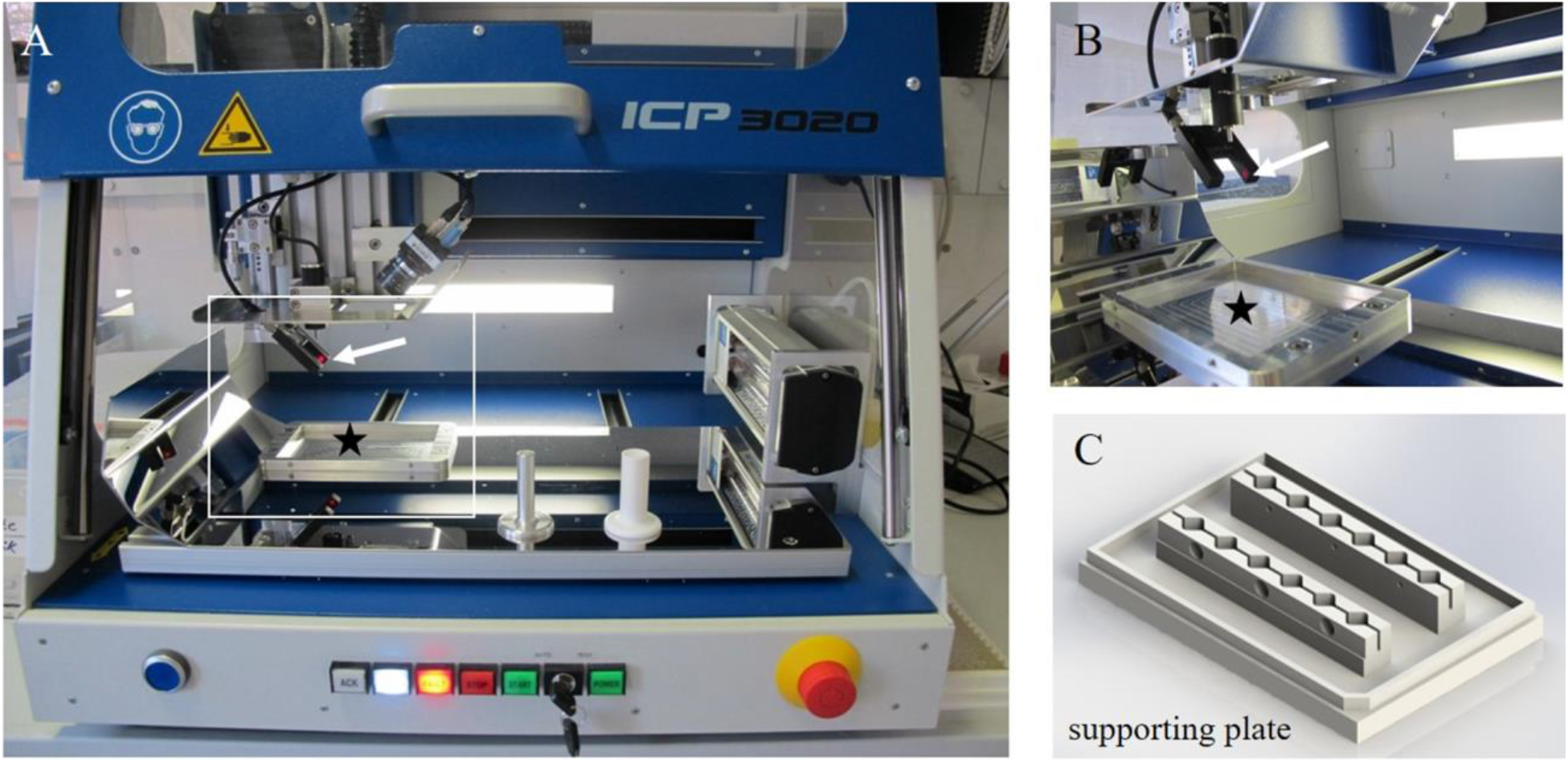
Artificial tissue cutter (ARTcut^®^): A) View of the computerized numerical controlled device equipped with B) optical barrier (white arrow indicates the laser light beam) and a C) supporting plate to fix the samples and place on the bottom of the machine (marked with black star).

After creation of full thickness or 1 mm chondral defects, the cartilage defects were filled with cell embedded collagen type I hydrogels (Figure 2). For subsequent *ex vivo* culture, osteochondral explants were transferred into custom-made culture platform [20] and cultured for 28 days with tissue specific media (Table 1) changed every 3-4 days. The *ex vivo* culture was carried out in a humidified atmosphere at 37°C and 5 % CO_2_ (BBD 6220 CO_2_ Incubator, Thermo Scientific™) either under normoxic (20 %) or physiological hypoxic (2 %) conditions.

**Table 1:**
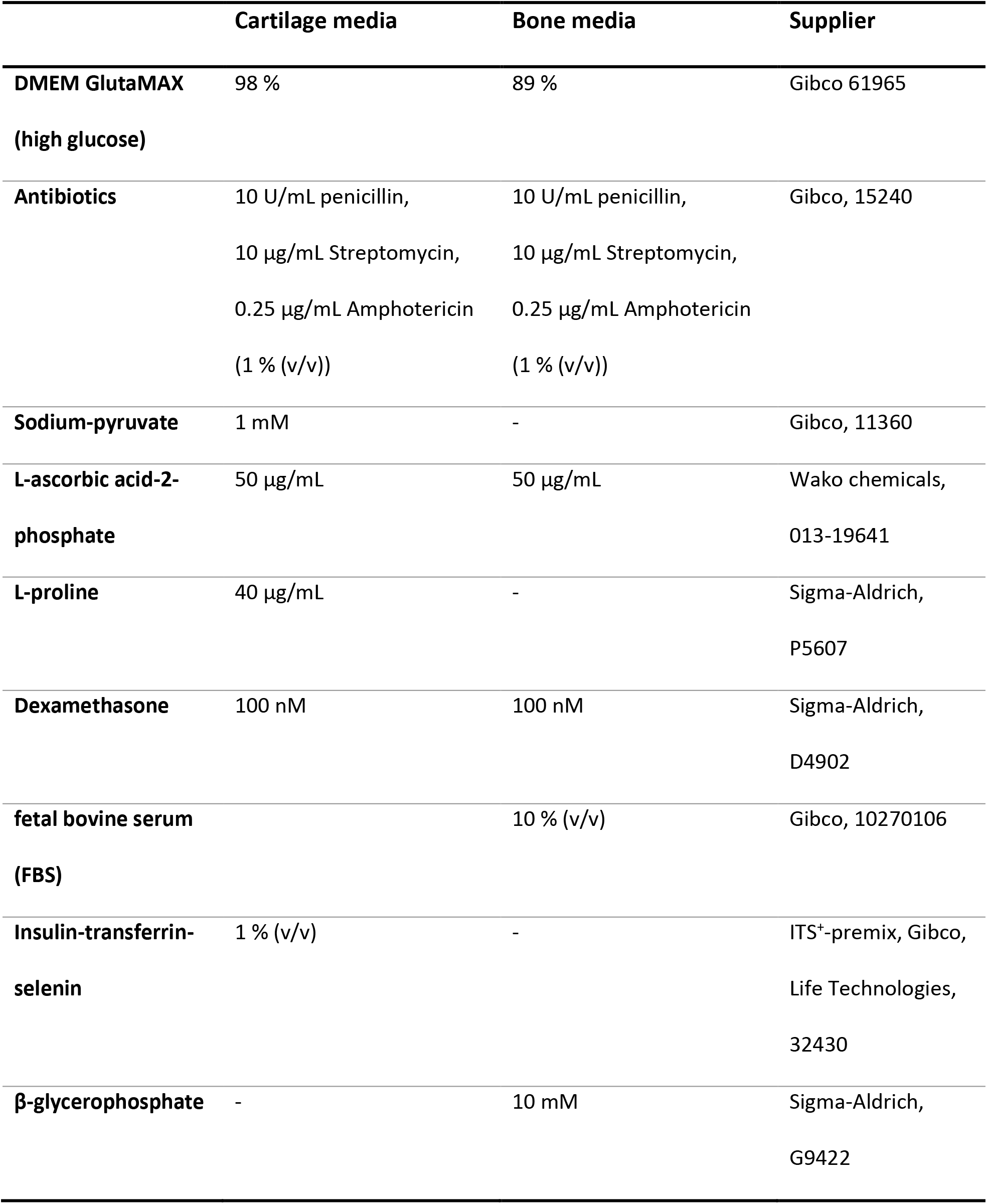
Composition of bone and cartilage media for *ex vivo* culture of osteochondral explants.

**Figure 2:**
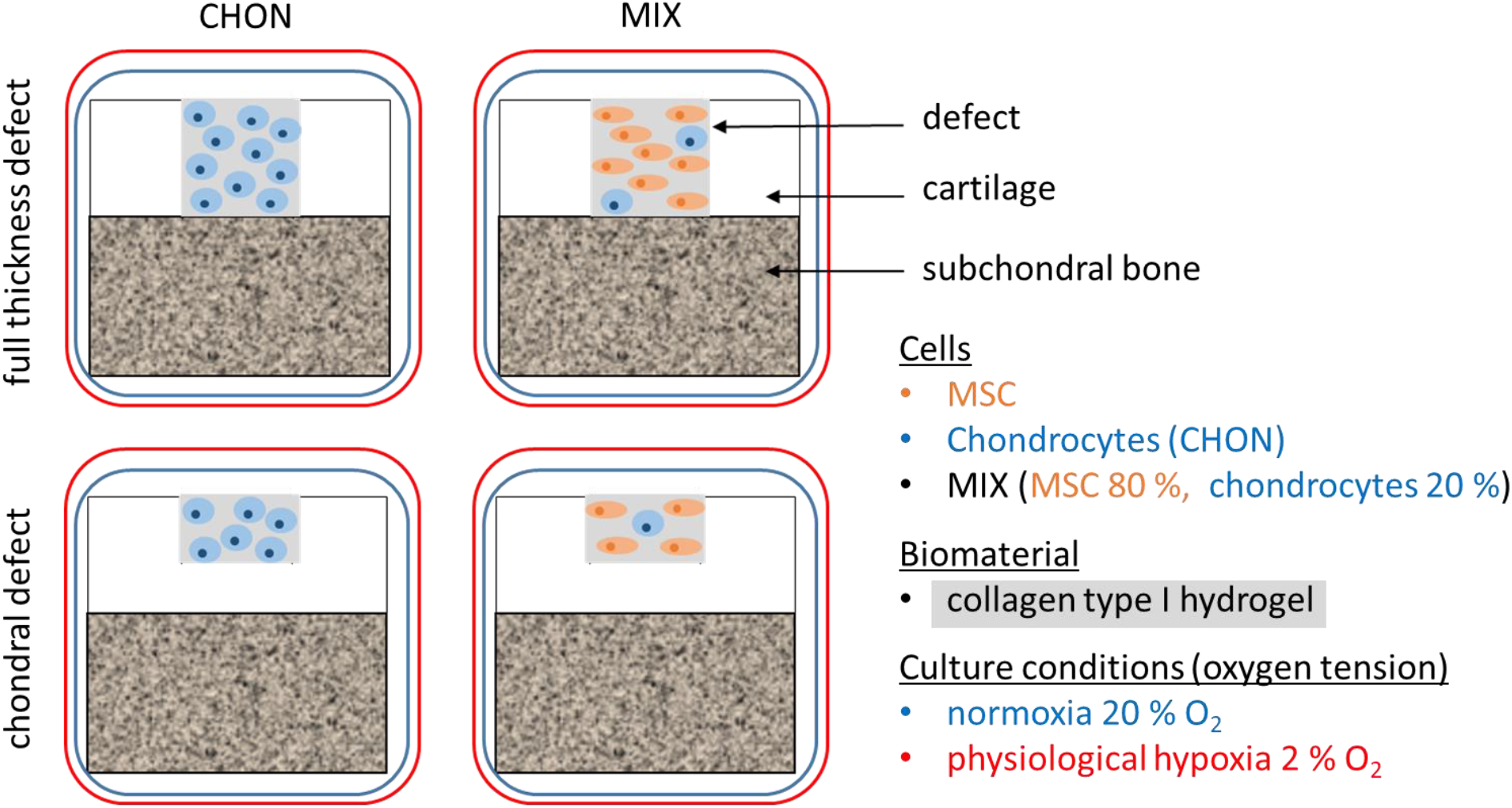
Schematic overview of all experimental groups: Full thickness and chondral defects of osteochondral explants were treated with collagen type I hydrogel (grey) either containing chondrocytes (CHON, blue) or MIX (80 % MSC [orange] and 20 % chondrocytes), implanted in chondral or full thickness defects and cultured *ex vivo* under normoxic (20 % O_2_, blue frame) or physiological hypoxic conditions (2 % O_2_, red frame).

### 2.2 Chondrocyte and mesenchymal stromal cells (MSC): isolation and *in vitro* expansion

Porcine chondrocytes were isolated from lateral condyles of 6–8-month-old domestic pigs by enzymatic digestion as previously described [20]. Chondrocytes were used at passage 0 – directly after isolation - for embedding in collagen type I hydrogel.

Porcine MSC were isolated from bone marrow aspirate (iliac crest) under the approval (reference number: 55.2 2532-2-256) of the District Government of Lower Franconia and the local animal welfare committee and performed according to the German Animal Welfare Act and the EU Directive 2010/63/EU according to the procedure described in chapter 1.2 in the supplementary I.

### 2.3 Cell embedding in collagen type I hydrogel

In a pilot study, cartilage defects were treated with collagen type I hydrogel (see chapter 1.3 in the supplementary for details). For implantation, two different cell laden constructs, as illustrated in Figure 2, were prepared and each cultured under normoxic and physiological hypoxic conditions. 1) CHON were prepared by embedding of chondrocytes in collagen type I hydrogel without expansion at a final density of 20 million cells/mL hydrogel volume; 2) MIX were prepared by mixing 80 % MSC with 20 % chondrocytes at same cell density. CHON and MIX implants were filled into the cartilage defect for gelation at 37°C.

### 2.4 Live-dead viability staining

To visualize a possible effect of defect creation process on cell viability of tissue samples, live-dead viability staining was performed (live-dead staining kit for mammalian cells, L3224, Invitrogen). Osteochondral explants were incubated with 4 μM Calcein AM and 2 μM ethidium homodimer-1 in DMEM HG and visualized with fluorescence microscope (494 nm/517 nm and 517 nm/617 nm wavelength) Keyence BZ-9000 (Biorevo). Living cells are stained by Calcein AM in green, dead cells are stained by ethidium homodimer-1 in red.

### 2.5 Histological evaluation

Osteochondral explants, harvested on day 0 and day 28, were washed with phosphate buffered saline, fixed for 24 h with 4 % formalin (Roti Histofix, Carl Roth P087.3) and processed with plastic embedding (Technovit T9100, Heraeus Kulzer, 66006735) as detailed described in the supplementary.

For immune-histological stainings, antigens were enzymatically retrieved depending on antibody (Table 2). Before incubation with primary antibody over night at 4°C, slides were blocked with 3 % (v/v) H_2_O_2_ (Carl Roth, 8070) and incubated with 5 % (w/v) bovine serum albumin. All stainings were visualized with Horseradish-peroxidase kit (DCS Innovative, Dako, K3468) following manufacturer’s instructions. Cell nuclei were counterstained with Mayer’s hematoxylin (Morphisto, 11895), rehydrated and mounted with Entellan (Merck, 1079610500). All washing steps between blocking or chemical incubation were performed with PBS supplemented with 0.5 % Tween-20 (VWR, 8.22184).

**Table 2:** List of antibodies for immune-histological stainings.

| Antibody | clone | Dilution and final antibody concentration | Antigen retrieval | Antibody supplier |
| --- | --- | --- | --- | --- |
| **Collagen type I** | EPR7785 | 1:1000; 0.875 µg/mL | pepsin & hyaluronidase | Abcam, ab138492 |
| **Collagen type II** | II-4C11 | 1:1000; 1 µg/mL | pepsin & hyaluronidase | MP Bio, 63171 |
| **Collagen type X** | COL-10 | 1:4000; 0.25 µg/mL | pronase | Sigma-Aldrich, C7974 |
| **Aggrecan** | EPR7785 | 1:2000; 0.5 µg/mL | pronase | Invitrogen, AHP0012 |

### 2.6 Histological scoring

Histological scoring for the evaluation of the immune histological stainings (collagen II, X and aggrecan) was performed. For in vitro tissue engineered samples the Bern score has been introduced, while cartilage tissue repair in clinical studies can be evaluated based on the international cartilage repair society (ICRS) score II [27]. Further, Chang et al. introduced a modified International cartilage repair society score by adding two new criteria, namely the visual assessment of collagen type I and II staining [28]. The inclusion of extracellular markers is an important step to better evaluate the composition of the repair matrix. Therefore, in this pilot study, the scoring category A (Bern-score), originally evaluating the uniformity and darkness of Safranin O-fast green stain, was translated to immune histological stainings of collagen type I, II, X and aggrecan) [29]. Stained sections (*n*=2 biological replicates) were evaluated blinded by three independent operators according to the criteria summarized in Table 3. Maximum possible score for the here reported study was 12 points.

**Table 3:**
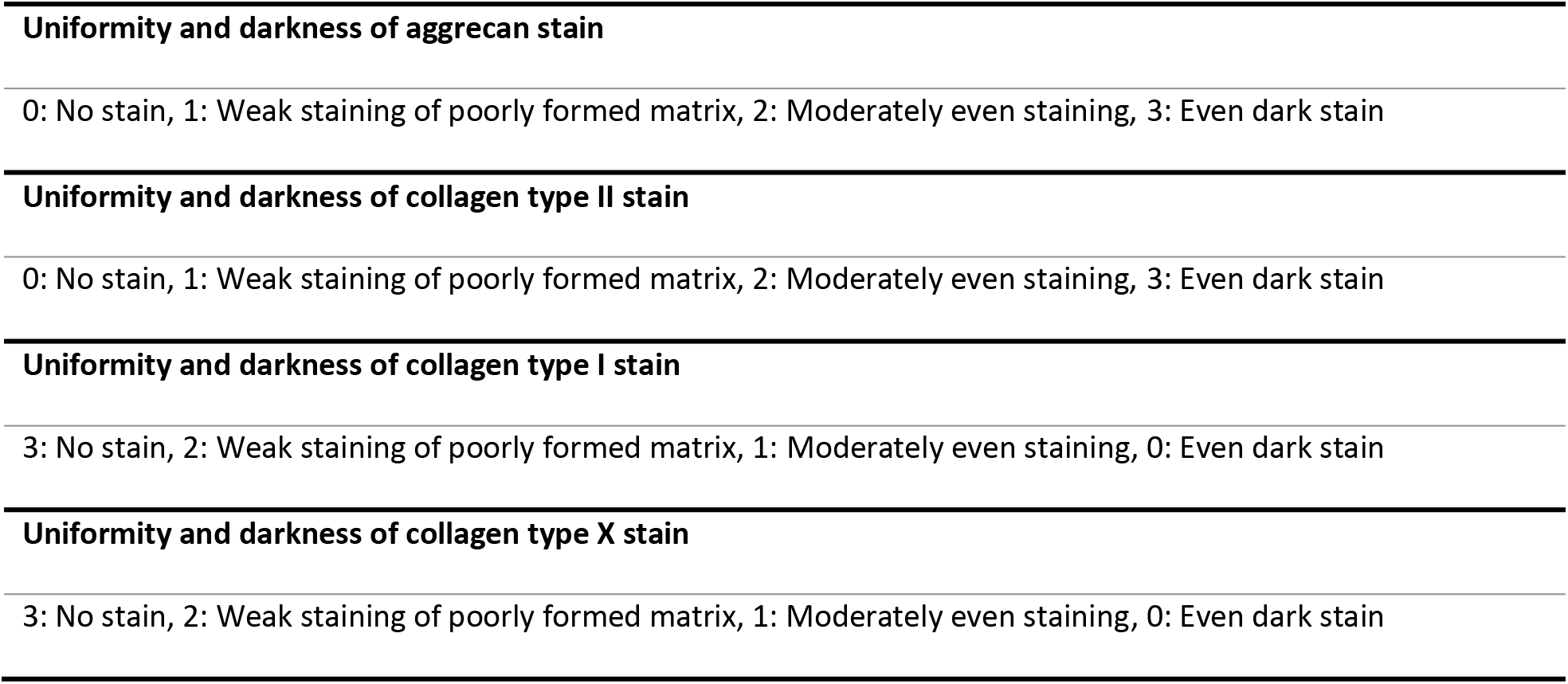
Criteria for evaluating immune histological stainings with a maximum of 12 points (0-3 scores for each criteria assessing the uniformity and darkness of aggrecan, collagen type I, II and X staining). Table adapted from Grogan et al. and Chang et al. [28, 29].

### 2.7 Statistics

Statistical analysis was performed with GraphPad Prism 6.07. Scoring values obtained from two independent experiments, each evaluated by three examinators were used for statistical analysis, resulting in *n* = 6 for each experimental group (Figure 2). Two groups differing in only one culture parameter (oxygen tension, defect depth and applied cell type) were directly compared. Following tests for outlier and Gaussian distribution, the data was either compared using an unpaired t-test (normally distributed values) or a Mann-Whitney test (not normally distributed values). A *p*-value <0.05 was considered statistically significant.

## 3 Results

### 3.1 Chondral defect creation with ARTcut^®^

ARTcut^®^ (Figure 1) represents an automated device for reproducible creation of chondral defects (1 mm in depth; Figure 3A). Main advantage of using the ARTcut^®^ is the ability to create defined chondral defects with a flat bottom to allow a tide and uniform contact with any solid pre-formed cylindrical implant, for example bioprinted constructs with flat bottom.

**Figure 3:**
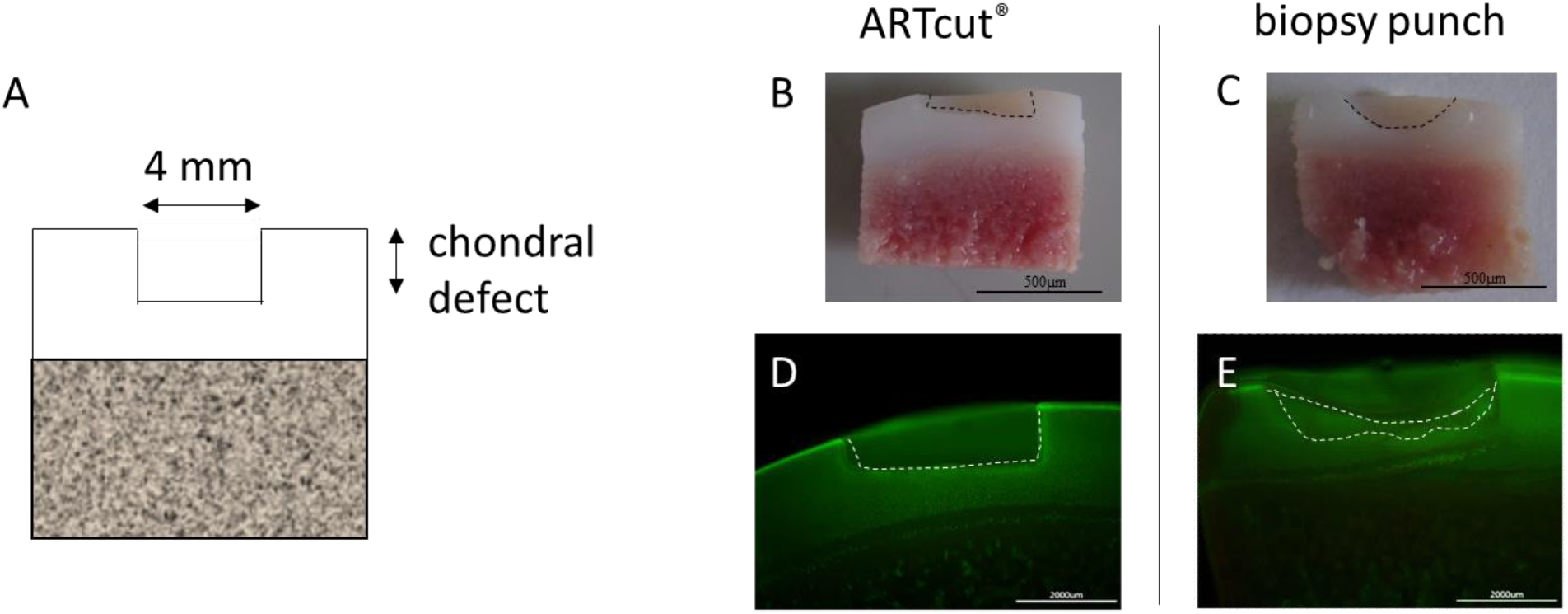
Osteochondral explant cross section with chondral defect. A) Schematic illustration of A) 4 mm chondral defect. B-E) Comparison of chondral defect created automated with ARTcut^®^ (B) and manually with biopsy punch (C) (scale bar 500 μm). D-E) Live-dead staining of explant cross section highlighting accurate borders of defect induced with ARTcut^®^ (D) and more irregular shaped geometry of manually induced defect (E) (scale bar 200 μm).

The laser beam to detect the surface of every single osteochondral explant is the unique feature of the ARTcut^®^ to define the onset of the drilling, thus all defects result with the same depth. Chondral cartilage defect induced with ARTcut^®^ appeared as a well-defined rectangular defect boundary in cross section (diameter: 4 mm, height: 1 mm) with flat bottom (Figure 3B). In contrast, the manually created defect (using biopsy punch) resulted in a half-round shape and required removing of residual cartilage tissue with a second tool (*e.g.,* a sharp spoon) shown in Figure 3C. Following, the accuracy of chondral defect geometry was much lower in manual procedure - using biopsy punch - compared to software – using automated ARTcut^®^.

The automated defect creation took less than 5 sec and thus heating of tissue due to friction and rotation during drilling was very unlikely. Microscopic images of live-dead stainings of osteochondral explants with 1 mm cartilage defects (diameter: 4 mm) confirmed no indications of cell death after tissue defect creation, neither with ARTcut^®^ nor with biopsy punch (Figure 3D-E).

For the creation of full thickness cartilage defects, the ARTcut^®^ was not required, since the bottom of the defect was given by the cartilage-bone interface. This two-tissue interface was easily separated using biopsy punch by pulling out the upper cartilage part.

### 3.2 Evaluation of cartilage treatments in the *ex vivo* model

Macroscopic images of all treatment and culture groups of the osteochondral *ex vivo* model after 28 days of culture are illustrated in Figure 4 A and B. The results of the pilot study demonstrated the feasibility and reproducibility of the *ex vivo* cartilage test system as a platform to compare cartilage repair strategies. Results of the adapted Bern scoring are shown in Figure 4 and Table 4 with scoring results of the co-culture (MIX) being close to the CHON treatment in chondral and full thickness defects. Representative sections of the scored samples are illustrated in Figure 5 and 6.

**Table 4:**
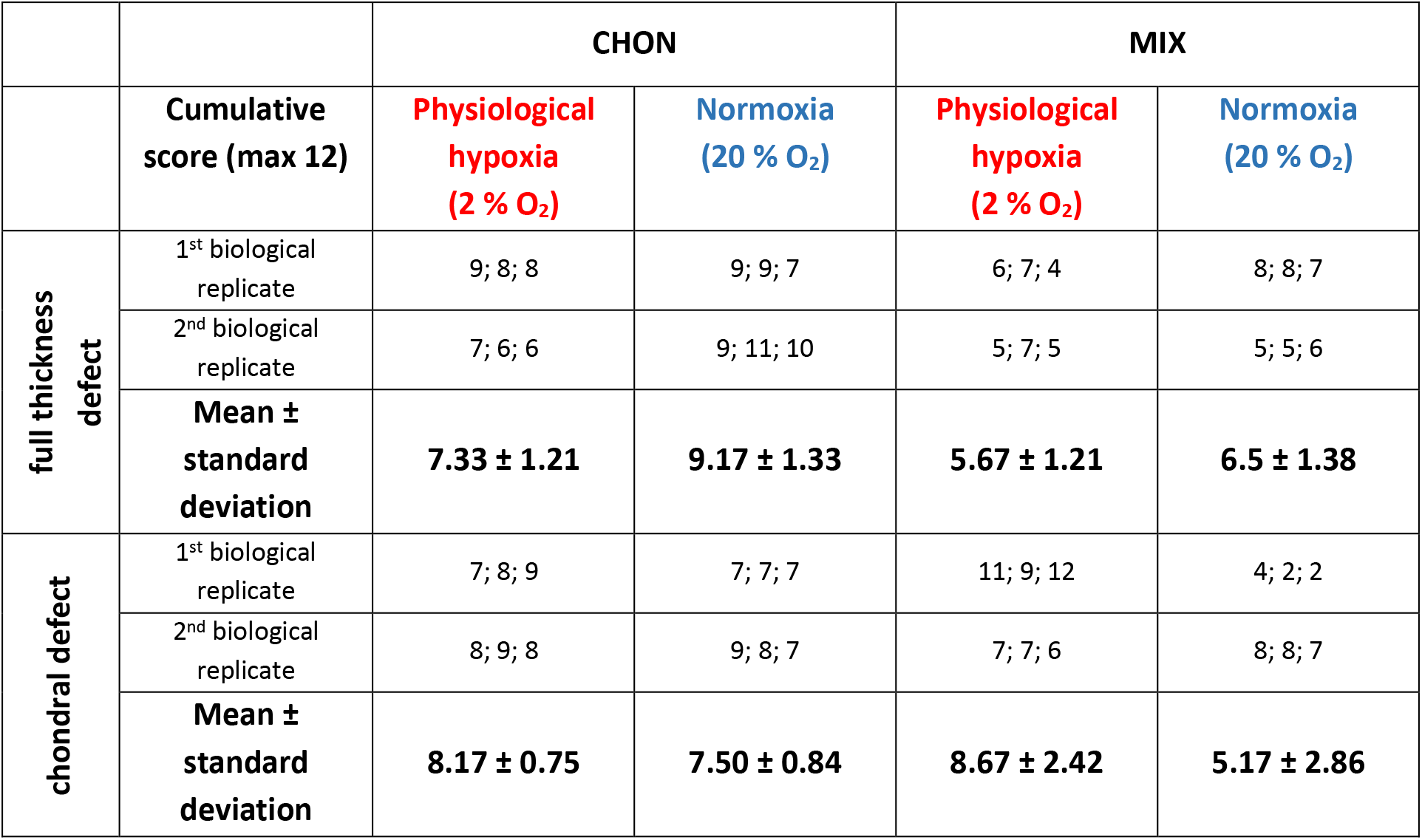
Scoring results to evaluate cartilage repair strategies in the ex vivo model. The table summarized the cumulative score of the 3-independent scorer (mean ± standard deviation with single scoring results of both biological replicate) for the CHON (100 % chondrocytes) and MIX (20 % chondrocytes, 80 % MSCs) treatment of the full thickness and chondral defects after 28 days culture. Values for samples cultured under physiological hypoxia (2 % O_2_) and normoxia (20 % O_2_) are displayed in separate columns.

**Figure 4:**
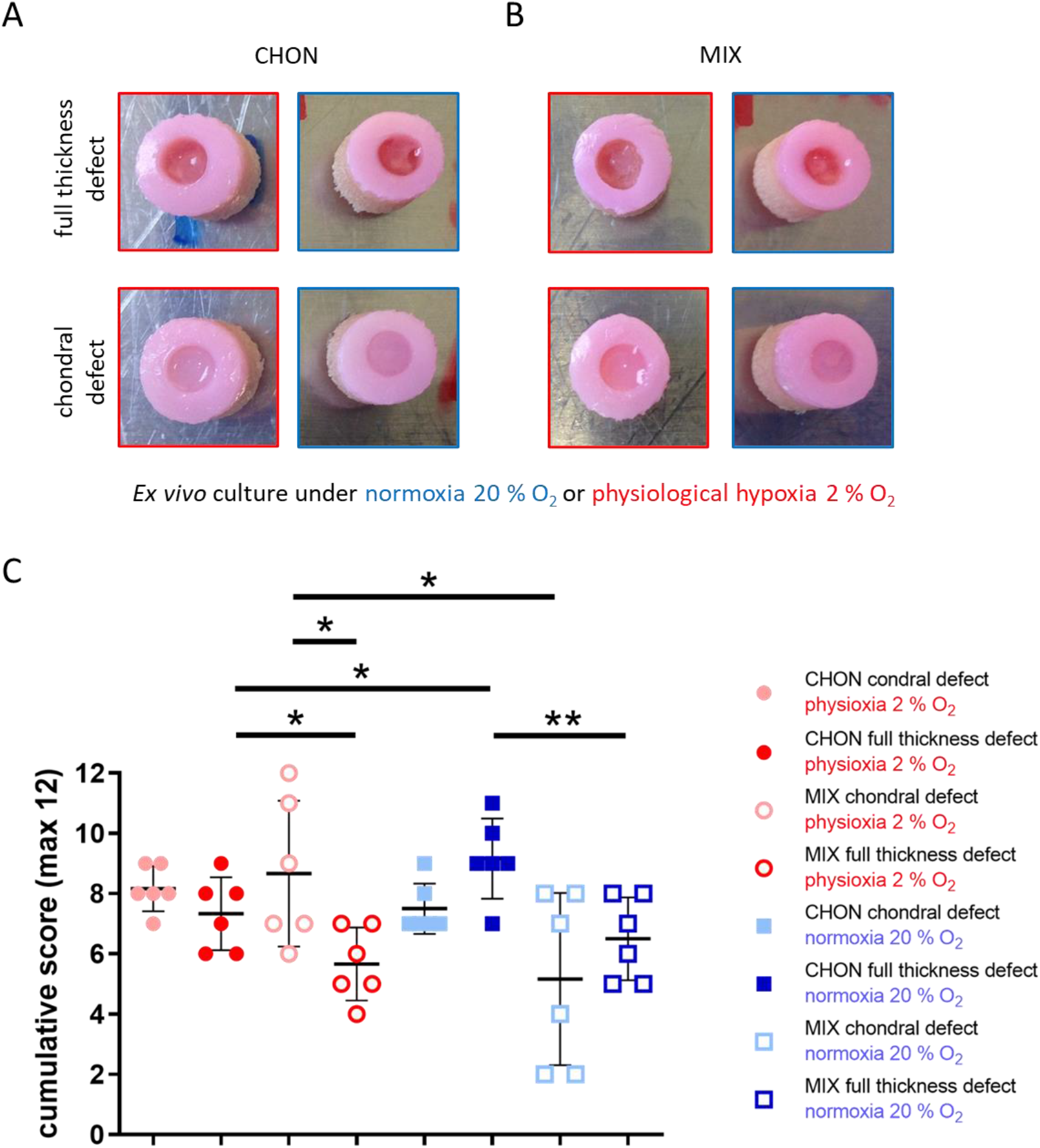
Evaluation of defect repair after 28 days *ex vivo* culture. Macroscopic images of full thickness and chondral defects (diameter 4 mm) treated with A) porcine chondrocytes (CHON) or B) co-culture of porcine MSC and CHON (MIX). Red frame indicates culture under physiological hypoxic, blue frame under normoxic oxygen tension. Macroscopically, there are no obvious differences comparing full thickness and chondral defects. C) Scoring evaluation of cartilage repair strategies in the *ex vivo* model (mean ± standard deviation with single scoring results as dots resp. listed values for 1^st^ and 2^nd^ biological replicate). Unpaired t-test or Mann-Whitney test * *p* < 0.05, ** *p* < 0.01.

**Figure 5:**
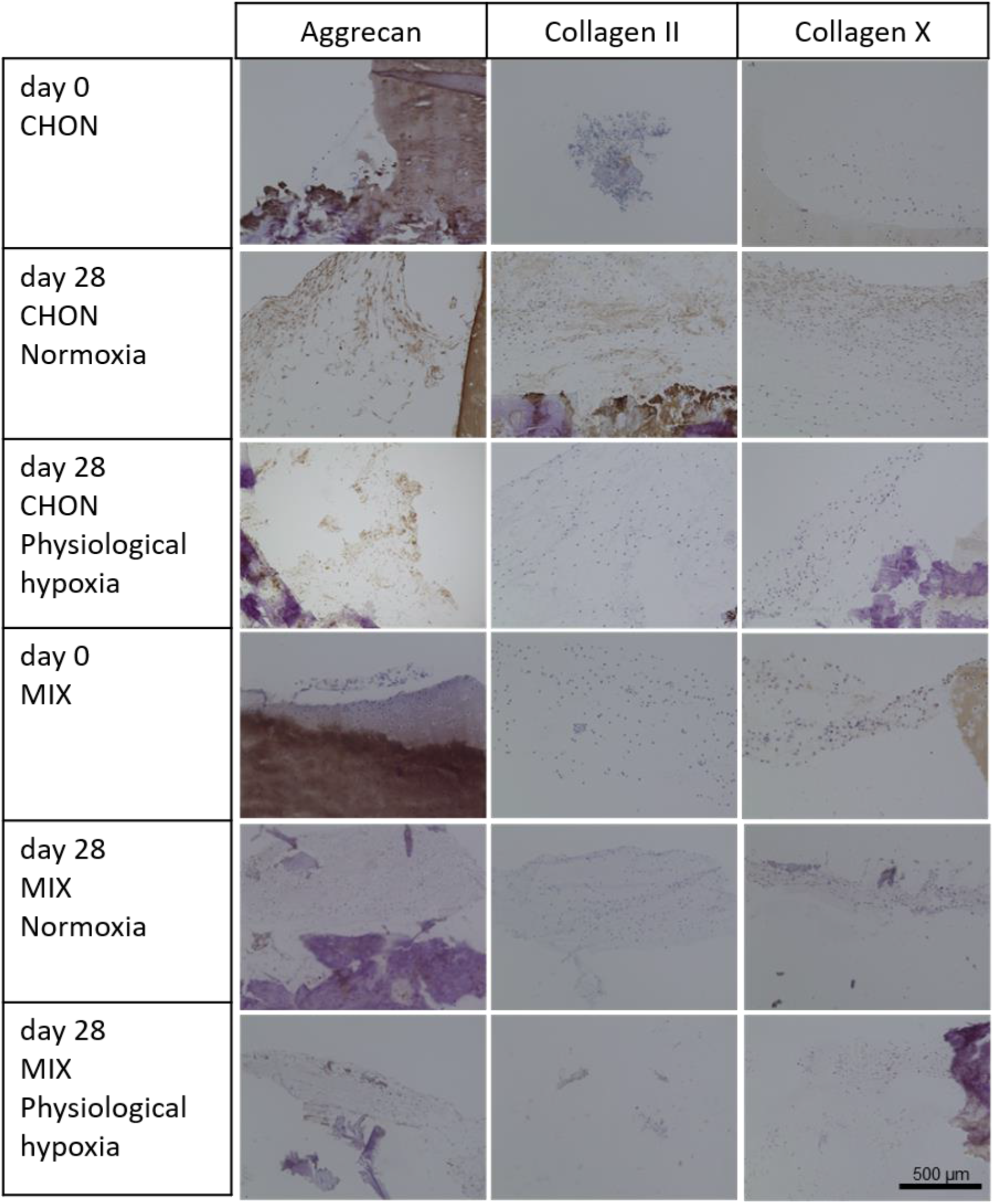
Chondral defects of the osteochondral defect model treated with either chondrocytes (CHON) or MSC-chondrocyte co-culture (MIX) embedded in a collagen hydrogel. Histological stainings of cross sections showing the defect area at day 0 and after 28 days of ex vivo culture under normoxic (20 % O_2_) or physiological hypoxic (2 % O_2_ conditions. Scale bar 500 μm.

**Figure 6:**
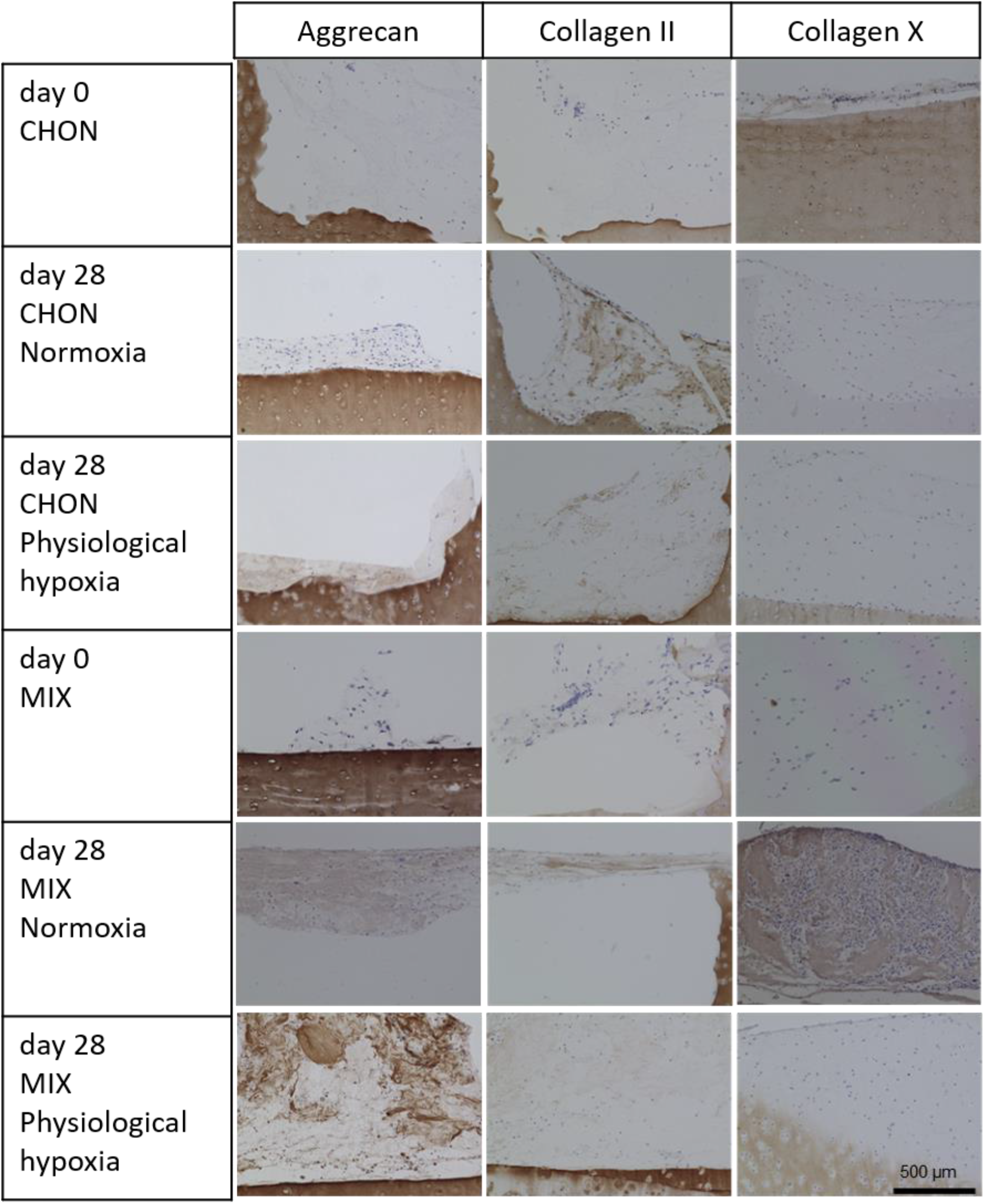
Full thickness cartilage defects of the osteochondral defect model treated with either chondrocytes (CHON) or MSC-chondrocyte co-culture (MIX) embedded in a collagen hydrogel. Histological stainings of cross sections showing the defect area at day 0 and after 28 days of ex vivo culture under normoxic (20 % O_2_) or physiological hypoxic (2 % O_2_) conditions. Scale bar 500 μm.

Under physiological hypoxic conditions, scoring results of defect repair after co-culture treatment (MIX) was higher (*p* = 0.0218) in chondral defect than in full thickness defects. Scoring values of CHON treatment in chondral and full thickness experimental groups did not differ much (n.s. *p* = 0.1828). At physiological hypoxic conditions the scoring value was higher for CHON treatment compared to MIX treatment in full thickness defects (*p* = 0.0384).

Under normoxic conditions, full thickness defects reached higher scoring values compared to the chondral defects.

Comparing the outcome of same defect depth and treatment group (cell type), but cultured under physiological hypoxic or normoxic conditions, the oxygen tension plays only a significant role in chondral defects using MIX (*p* = 0.0451, higher scoring values under physiological hypoxic conditions) and in full thickness defects treated with CHON (*p* = 0.0316, lower scoring values under physiological hypoxic conditions).

Overall, treatment of the two defect depths with MIX showed slightly reduced scoring values compared to the CHON treatment group. Only exception is MIX chondral (physiological hypoxia) with a slightly higher value as the corresponding one for CHON. The results of the MIX treatment groups also tend to spread more than the CHON treatment group independently of defect depth and oxygen tension.

## 4 Discussion

The choice of optimal treatment approach for cartilage repair is dependent on the size and depth of the cartilage defect and thus the severity of pathological condition.

To mimic these different pathological defect scenarios, the implementation of the ARTcut^®^ allowed to overcome the low reproducibility of manual induced chondral defects. So far, there is no other automated defect creation device published that has comparable features, namely possibility to select different drilling parameters within one run and the application of one device for a wide range of soft and hard tissues or tissue equivalents, as it is demonstrated with the ARTcut^®^. The induction of defects in osteochondral explants allows to compare different cell-based treatment approaches by implantation of cell loaded materials into the defects or to study the effect of defect depth on repair in an *ex vivo* model. The software-controlled drilling parameters allow for standardized and reproducible defect creation in other soft and hard tissues including bone tissue as well. *In vitro* as well as *ex vivo* studies require sample procession at high throughput rates under a negative microbial environment. ARTcut^®^ device addresses these demands, resulting in higher methodological accuracy and reproducibility, compared to manual procedure using biopsy punch.

Current treatments of small or large cartilage defects mainly aim on pain and symptom relief rather than on functional tissue repair [30]. From a scientific point of view, there is way for improvement of cartilage defects, especially on long term results. Material-assisted as well as cell-based therapies, including (M-) ACI, AMIC and microfracture, tend to result in formation of mechanical inferior fibrocartilage in long term follow up [30–32]. Moreover, the treatment of large defects with autologous cells are limited due to the high number of chondrocytes needed for the current techniques. Therefore, in this laboratory-controlled study, an *ex vivo* osteochondral defect model was modified to screen cell-based approaches to treat chondral and full thickness cartilage defects. Different cell types, namely chondrocytes and MSCs, were embedded in collagen type I hydrogel (CHON or MIX) and implanted in trauma induced chondral and full thickness defects. A tendency of a stimulative and beneficial effect of MIX treatment (20 % CHON and 80 % MSCs) was observed regarding cartilaginous matrix formation with the main advantage to reduce the overall number of CHON used for traditional ACI treatment. It has been demonstrated in previous studies that the cells in cartilage and subchondral bone remained metabolic active up to 56 days of *ex vivo* culture [20].

Promising results regarding cartilage matrix production were obtained in this pilot study comparing CHON and MIX in chondral and full thickness defects. The most pronounced increase in proteoglycan content normalized to DNA amount resulted for the MIX treatment in both conditions, chondral and full thickness defect, cultured under physiological hypoxia (Supplementary Figure S1). The MIX treatment reduces the amount of autologous cartilage tissue needed for chondrocyte isolation, since chondrocytes represented only 20 % of the total cell number in our model. Several *in vitro* studies and *in silico* models have shown that chondrocytes in co-culture with MSCs increase the chondrogenic differentiation potential [18, 19, 33, 34]. Scalzone *et al.* reported of an enhanced chondrocyte activity measured by proteoglycan and collagen type II production after co-culture with MSC compared to chondrocyte monoculture both embedded in chitosan based hydrogels [19].

One advantage of the co-culture approach – when clinically applied – would be the one-step procedure, addressing the surgical intervention of the knee joint, for the treatment of the cartilage defect: The reduced number of chondrocytes in the MIX treatment can be isolated intra-operatively without expansion and thus reduced the risk of chondrocyte dedifferentiation [35]. Further, the required MSCs, *e.g.,* can be harvested independently form the knee surgery from bone marrow aspirate in a minimal invasive procedure with subsequent expansion prior to the cartilage repair procedure to ensure a sufficient number of MSCs.

Considering the absence of TGF-β in the culture media, which is known to drive MSC chondrogenesis and matrix production [36], the here reported cartilage repair solely derives from the stimulating effect of the surrounding osteochondral tissue, cell-cell signaling of the cell laden implants and the culture conditions. Therefore, the overall matrix deposition is rather low due to missing stimuli of exogenous growth factors in the here described model. Results of a similar *ex vivo* model, based on horse and bovine osteochondral explants, suggested that the presence of the osteochondral tissue in the explants increase cartilage-like matrix deposition compared to free swelling conditions of cell seeded material only [21, 22].

Beside the cell source, the influence of oxygen tension (normoxia 20 % O_2_, physiological hypoxia 2 % O_2_) was studied during the *ex vivo* culture on chondrogenic matrix deposition, resulting in different outcomes between the treatment groups in chondral and full thickness defects. Based on the histological scoring, the results of the pilot study showed that the oxygen tension seems to only plays a role in chondral defects - marked by elevated matrix production. In contrast, full thickness defects showed different response in cartilage-like matrix deposition related to the treatment approach: While higher scoring values were reached with CHON treatment under normoxic conditions, the pMIX treatment did not show differences comparing physiological hypoxic and normoxic conditions. Physiological hypoxic conditions resulted in an increase in cartilaginous matrix deposition in chondral defects compared to full thickness defects independently of the used cell type (CHON, MIX). Once the defect reaches the cartilage-bone interface, the cells in the implant receive stimuli from the subchondral bone that seem to counteract with the hypoxic induced stimuli in the here presented model. One explanation of the differential cell response to oxygen tension in chondral and full thickness defects may originate from the variation in oxygen tension present in the human body. Chondrocytes are exposed to lower oxygen tension in avascular cartilage (2-5 %) [37], synovial fluid and the synovial capsule (6.5-9.0 %) [25, 38], with an increasing oxygen level in the bone marrow of subchondral bone (>7 %) [26, 39] and a maximum of 12 % in arterial blood [40]. Once the cartilage defect progresses to the exposure of the subchondral bone in patients, chondrocytes within the defect are exposed to higher levels of oxygen present in the bone supplied by vascular invasion [41]. It has been shown that low oxygen inhibits the degradation of hypoxia induced factors (HIF) [42]. HIFs are described to be essential for maintaining CHON homeostasis and extracellular matrix synthesis and activate the transcription of genes [43].

While the here presented results of the pilot study are consistent, traceable and in line with literature, there are still limitation in the current experimental setup: Due to limitations in availability of human healthy osteochondral tissue – absence of any osteoarthritic phenotype – the authors decided to use porcine tissue instead, as introduced by Schwab *et al.* [20]. The performance of an extensive comparative *ex vivo* study comprising three parameters – namely 1) defect depth (chondral vs. full thickness), 2) different cell types for defect filling and treatment (100 % chondrocytes vs. 20 % chondrocytes and 80 % MSCs), and 3) oxygen tension (physiological hypoxia vs. normoxia) requires tissue material in the required quantity and of similar quality. Due to the well-known donor issues associated with donor variability of human tissue and the already mentioned limited availability of healthy human osteochondral tissue, this could not be achieved for human tissue [44, 45]. In contrast, the use of porcine tissue isolated from one pig population (similar age; grown up under equal conditions) minimizes the donor variation and represents a tissue source for healthy and non-degenerated tissues with higher reproducibility. To proof the findings and the trend of the here presented study and to show clear differences between treatment groups, this experimental set-up needs to be repeated with more biological replicates.

The results obtained in this pilot study indicate that the surrounding tissue of the osteochondral explants plays a crucial role in the defect repair, addressing chondral or full thickness defects, and that the oxygen tension additionally stimulates the outcome in a defect depth dependent way. However, the exact mechanism controlling success or failure of a treatment approach are not fully understood and require further research. Despite the advantage of the here reported model, the mimicry of the complex 3D environment present in the knee joint with the additional ability to reproducibly create (trauma induced) defects, this model has its potential in pre-screening several biomaterials or treatment approaches, but best candidates still require further pre-clinical and clinical testing to confirm obtained results.

To conclude, in this *ex vivo* study two cell-based approaches (CHON vs. MIX embedded in collagen type I hydrogel) for the treatment of chondral and full thickness defects, cultured under normoxic and physiological hypoxic conditions, were compared on their cartilage repair potential. Independently of defect depth, the scoring results of the co-culture of chondrocytes and MSC (MIX) are close to those of CHON treatment. Hence, usage of MIX could be a promising approach to reduce the number of chondrocytes to one fifth and thus the amount of tissue for harvest and accompanied side morbidities for the patient. Further, in this experimental set up the oxygen tension showed an influence in a defect depth dependent way that has not been reported in literature before. The here presented advanced osteochondral cartilage defect model provides a promising platform to deeper understand the underlying mechanism.

## Acknowledgements

The authors thank Andreas Diegeler (Fraunhofer ISC, Bronnbach) for technical support of the ARTcut^®^. Thanks also to Sebastian Naczenski for establishing the protocol for wounding of osteochondral explants using ARTcut^®^ device. Finally, we would like to thank Heike Walles for her support. We acknowledge the authors of the thesis (Andrea Schwab and Alexa Buss) for the permission to use the data in this manuscript. Parts of the illustrations were produced in the light of a doctoral thesis (figure 1 and 3, with approval from Schwab 2017, German copyright law) [46] and a doctoral thesis (figure 5 and 6, with approval from Buss 2021) [47].

## Author contribution

Conception and design of the study: Franziska Ehlicke and Andrea Schwab; Acquisition of data: Alexa Buss and Andrea Schwab; Analysis and interpretation of data: Andrea Schwab, Alexa Buss, Franziska Ehlicke and Oliver Pullig; Manuscript drafting: Andrea Schwab and Alexa Buss; Revision of the manuscript: Franziska Ehlicke and Oliver Pullig; Final approval: Franziska Ehlicke, Andrea Schwab, Alexa Buss and Oliver Pullig; Funding: Franziska Ehlicke and Alexa Buss.

## Role of funding Sources

This work was financially supported by the European Union Seventh Framework Programme (FP7/2007-2013) under grant agreement no 309962 (HydroZONES).A.Buss was supported by a medical scholarship of the German Excellence Initiative to the Graduate School of Life Sciences, University of Würzburg. This publication was supported by the Open Access Publication Fund of the University of Wuerzburg.

## Conflict of interest statement

The authors declare no conflict of interest.

## Ethics approval and consent to participate

Animal experiment was approved (reference number: 55.2 2532-2-256) by the District Government of Lower Franconia and the local animal welfare committee and performed according to the German Animal Welfare Act and the EU Directive 2010/63/EU. Following heparinization of the pig, porcine MSCs were obtained by bone marrow aspiration.

## Notes

### Competing Interest Statement

The authors have declared no competing interest.

### Summary of Updates

Clarification of the use of the modified BERN score were added to the methodology. The histological stainings of the scored sections have been added (a selectin of images) to the manuscript. GAG Data was added to the supplementary to compare with scoring results.

