## Supplementary Information for "*Ex vivo* osteochondral test system with control over cartilage defect depth – a pilot study to investigate the effect of oxygen tension and chondrocyte-based treatments in chondral and full thickness defects in an organ model"

### 1.1 Cartilage defect creation: Artificial tissue cutter

For standardized wounding of osteochondral tissues, we developed a support plate allowing simultaneous fixation of maximum 12 osteochondral cylinders (Figure 1C). The wounding procedure (Figure 1) is divided into four steps: change of millers with subsequent calibration run, parameter settings, device disinfection and wounding.

Due to the intended wound size and geometry (4 mm diameter), an appropriate miller (Dieter Schmid fine tools, Germany, 360014) was selected and mounted in the milling head. The wounding process is controlled by PALPC software V2.01, using the following parameter: drilling depth (1 mm) and drilling speed (80 Hz). The movable milling machine along x-, y- and z-axis allows for wounding of various samples within one run, defined by the number of fixated osteochondral cylinders in support plate.

### 1.2 Isolation of mesenchymal stromal cells

Following gradient centrifugation using Ficoll-Paque (GE Healthcare Bio-Sciences, 17-5442-03), mononuclear cell fraction was expanded in DMEM HG medium (GlutaMax, Gibco 61965) supplemented with 1 ng/mL FGF-2 (R&D Systems, 233-FB-025), 10 % FCS (Gibco, 10270106) and 1 % Antibiotic-Antimycotic (10 U/mL penicillin, 10 µg/mL Streptomycin, 0.25 µg/mL Amphotericin, Gibco, 15240) in humidified atmosphere (37°C, 5 % CO2). Media change was performed every 3-4 days. MSC were used at passage 2-3 for embedding in collagen type I hydrogel.

### 1.3 Collagen type I hydrogel: hydrogel preparation and cell embedding

Collagen was isolated from rat tails (Wistar IGS Rat; Charles River Laboratories), dissolved in 0.1 % (v/v) acetic acid to a concentration of 4.5 mg/mL, and mixed with half volume of gel neutralizing solution (GNS) to achieve a final concentration of 3 mg/mL. GNS consists of 2x DMEM high glucose (DMEM, PAA Laboratories, G0001,3010), 0.09 M HEPES (Sigma, H3375), 3 % FCS (Bio&Sell, FCS.ADD.0500), 0.05 mg/mL chondroitin sulphate (Sigma,

C4384) with pH value adjusted to pH 8.5. Cell embedding was realized by mixing the porcine chondrocytes and/or MSCs in GNS, resulting in a final total cell concentration of 20 Mio/mL hydrogel. Gelation was performed via incubation of GNS collagen solution mixture for 20 min at 37°C under humidified atmosphere.

### 1.4 Plastic embedding

Osteochondral explants at day 0 and day 28 were washed with phosphate buffered saline, and subsequently fixed for 24 h with 4 % formalin. Samples were dehydrated in series of alcohol (ethanol 70 %, 80 % and 96 %, isopropanol I-III, xylene I-II) for 3 h each, incubated in pre-infiltration solution I-III for 24 h each and 72 h with infiltration solution (Technovit T9100, Heraeus Kulzer) according to manufacturer`s instructions. For pre-infiltraton step III and infiltration T9100 basis solution was destabilized through aluminum oxide filled column to allow immune-histological stainings. Polymerization of samples was performed for at least 24 h in Teflon molds after degassing in fridge. Samples were cut in 4 µm slices with rotation microtome (Leica RM 2255), mounted on object slides with 70 % ethanol, covered with PE foil and dried overnight at 60°C for attachment of slices on glass slide. Prior to histological staining, slides were deplastified with xylene (2 times 20 min), methoxy-ethyl-acetate (2 times 20 min), acetone (2 times 10 min) and washed with deionized water.

### 1.5 Quantification of proteoglycan and DNA content

For the quantification of proteoglycan (GAG) and DNA content, the cell laden hydrogels were digested with Papain digestion buffer (10mM phosphate buffer, 5mM L-Cystein, 500 mM EDTA, 140 µg/ml papain) at 50°C for 24h. GAG quantification was performed using the Blyscan™ Glycosaminoglycan Assay (biocolor, B1000) following manufacturer instruction. DNA content was analyzed using the Quant-iT™ PicoGreen® dsDNA Assay (Molecular probes Invitrogen detection Technologies, 7589) according to manufacturer instruction. Absorbance and fluorescence readout was done using the TECAN sunrise (TECAN) spectrophotometer. DNA and GAG content was calculated based on the respective standards. Figure S1 illustrates the results of the GAG/DNA data for the different treatment groups in the chondral and full thickness defects.

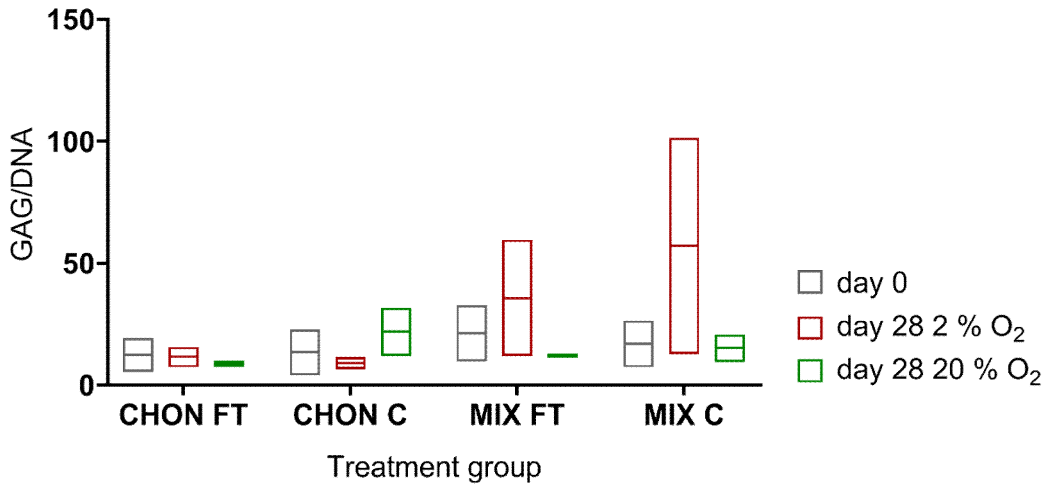


*Figure S1: Quantification of GAG/DNA in the hydrogel samples at day 0 and after 28 days of culture in the osteochondral defect model. Chondrocytes (CHON) or MSC-chondrocyte co-culture (MIX) were embedded in a collagen hydrogel and implanted in a chondral (C) or full thickness (FT) defect. MIX treatment showed an increase in GAG/DNA in both treatment groups after culture under physiological hypoxia (2 % $O_2$). CHON group resulted in increase in GAG/DNA only in chondral group cultured under normoxia (20 % $O_2$). Box plots illustrate min and max value with the mean value shown as line. FT: full thickness defect, C: chondral defect.*
